# Target tracking behaviour of the praying mantis *Sphrodromantis lineola* (Linnaeus) is driven by looming-type motion-detectors

**DOI:** 10.1101/2020.06.02.129684

**Authors:** F. Claire Rind, Lisa Jones, Ghaith Tarawneh, Jenny F. M. Read

**Affiliations:** Biosciences Institute, Newcastle University, NE2 4HH, Newcastle upon Tyne, UK

**Keywords:** Praying mantis, target, motion-detector, lateral-inhibition, looming

## Abstract

We designed visual stimuli to characterise the motion-detectors that underlie target tracking behaviour in the mantis. The first was a small, moving, stripy, bug-like target, made by opening a moving, Gabor-filtered window onto an extended, moving, sinewave pattern. The mantis tracked this bug-like target, but the likelihood of tracking the bug depended only on the temporal frequency of its motion. In contrast, optomotor responses to the extended moving sinewave pattern alone depended on both spatial and temporal frequency of the pattern, as expected from classical, correlation-based motion-detectors. In another experiment, we used small moving objects that were made up of chequerboard patterns of randomly arranged dark squares, and found objects with smaller sized chequers were tracked relatively less. Response suppression like this, when the internal detail of an object increases, suggests the presence of lateral inhibition between inputs to the motion-detectors for target tracking. Finally, wide-field motion of a chequerboard background near the target, balanced so no optomotor responses were evoked, suppressed tracking proportionally both to the nearness of the background to the target and to the size its dark chequered squares. Backgrounds with smaller sized squares produced more suppression. This effect has been used as a demonstration of lateral inhibition in detectors for looming-motion and makes their response greatest to an expanding outer edge, an image produced by an approaching object. Our findings point to a new role for a looming-type motion-detector in mantis target tracking. We also discuss the suitability of several large lobula-complex neurons for this role.

**Summary Statement:** Lateral inhibition shown by motion-detectors underlying target tracking by the praying mantis *Sphrodromantis lineola* (Linnaeus).

## Introduction

An animal needs to process many different motion cues in the environment and react appropriately to each. For example, insects show optomotor responses, mediated by classical motion-detectors that respond to spatiotemporally correlated changes in luminance, to correct unintentional motion of large parts of their visual field over their eyes (Reichardt and Guo, 1986) or, they can show tracking responses to motion of a small dark target object (Egelhaaf, 1985; Keleş and Frye, 2017; Nityananda et al., 2018), or, if the object’s image suddenly increases, it could indicate an approaching predator, and it can trigger an escape reaction (Fotowat et al., 2011; Rind and Simmons, 1999; Santer et al., 2006). The term target is used for small, usually dark objects that move independently from the background (Gonzalez-Bellido et al., 2016) and these targets can either be tracked as prey, as a mate, or can be avoided. Moving striped patterns, such as sinusoidally-modulated luminance-gratings are a good way of studying the sensitivity of any visual system to the luminance changes accompanying image motion, because in them the luminance changes regularly in time and in space over the eye and the underlying mathematical operations can be inferred, as long as the operations involve linear interactions. This assumption is most likely to be met with wide field motion (WFM) and the classical correlation based motion detecting pathways that underlie it (Kulikowski and Tolhurst, 1973; Reichardt and Poggio, 1979; Thompson, 1982). However, over limited regions of space, sinusoidally modulated luminance gratings can also reveal if the same mathematical operations underlie small-object movement detecting pathways. Sensitivity to these moving sinusoidally-modulated luminance-gratings can be used as a tool to reveal not only the visual system’s resolving powers in space; but also its temporal resolving powers (Kulikowski and Tolhurst, 1973; O’Carroll et al., 1997; Reichardt and Poggio, 1979; Thompson, 1982). Both neurophysiological and behavioural studies have shown that many insects’ contrast sensitivity to WFM is dependent upon the spatial and temporal frequency of a moving sinewave grating, demonstrating the insect WF motion detection system is tuned to a combination of spatial and temporal properties of a visual stimulus rather than a unique pattern velocity (Dvorak et al., 1980; Hausen and Egelhaaf, 1989; Lawson and Srinivasan, 2020; Pick and Buchner, 1979; Reichardt and Guo, 1986; Reichardt and Wenking, 1969; Straw et al., 2008). Different species of insect have evolved sensitivity to particular spatial and temporal frequencies which match their behavioural ecology (Lawson and Srinivasan, 2020; Nityananda et al., 2015; O’Carroll et al., 1997). Fast moving insects, such as flies and bumblebees, have evolved motion detection systems that are sensitive to spatial and temporal frequency combinations which represent high velocities (O’Carroll et al., 1997). In contrast, insects such as hoverflies, which are stationary when hovering but also make quick flights, have a sensitivity to both high and low velocities (O’Carroll et al., 1997). A recent study into the contrast sensitivity of the mantis *Sphodromantis lineola* to WFM has shown that the optomotor pathway in this insect predator is tuned to a broad range of spatial and temporal frequencies representing image motion speeds from 20 to 500 degrees per second (Nityananda et al., 2015).

Praying mantises are opportunistic predators using their spiny raptorial forelegs to catch a wide range of prey (Corrette, 1990; Prete, 1992; Prete, 1993). A mantis primarily uses its visual system to detect, and track prey before eliciting a predatory strike to capture them (Corrette, 1990; Rossel, 1980; Rossel, 1983; Yamawaki, 2000; Yamawaki and Toh, 2003). Sphodromantis lineola tracks square black targets ranging in size from 10° to 48° and strikes at targets subtending 12° x 12° on their eyes (Nityananda et al., 2019; Nityananda et al., 2018; Nityananda et al., 2016; Prete, 1993; Prete, 1999; Prete et al., 2002). As the distance of the target from the mantis increases beyond 25 mm, the likelihood of a strike decreases (Nityananda et al., 2016). It is one of the only animals known to determine that a target is in range using motion stereopsis, matching the disparate positions, or disparity, of small-field motion on the two eyes (Nityananda et al., 2018). Before this, the target must be brought and maintained within reach and this is achieved by tracking. We devised visual stimuli to determine the properties of the motion-detectors used to track a moving target. The stimuli allowed us to discriminate between correlation-type motion-detectors, like those of the optomotor pathway, (Keleş and Frye, 2017), and, looming-type motion-detectors excited by jittery motion of a small target, but responding best when the object approaches (Simmons and Rind, 1992; Simmons and Rind, 1997).

## Methods

### Experimental Model and Subject Details

Experiments were carried out on 20 adult African lined mantises (*Sphodromantis lineola*), acquired from a UK breeder. They were housed individually in plastic boxes (170 mm length, 170 mm width, and 190 mm height) with holes in the lid for ventilation. The housing facility was maintained at 25°C with a 12 hour light/dark cycle and the boxes were regularly misted with water to raise the humidity. They were fed one medium-sized field cricket (18 – 25 mm) twice per week.

### Experimental set-up

Mantises were individually positioned in front of a CRT monitor upon which visual stimuli were presented. Subjects were positioned underneath a Perspex perch (50 × 50 mm), which was clamped 60 mm away from a CRT screen (Phillips 107B3 colour CRT monitor 330 mm × 248 mm, refresh rate 85Hz). The monitor had a resolution of 1280 × 960 pixels, with a pixel density of 3.875 pixels mm^-1^. The monitor was gamma corrected using a Minolta LS-100 photometer. The viewing distance of the screen for each mantis was 60 mm with a visual angle of 140°horizontally and 128°vertically. A wooden box (66cm L × 530 mm W × 600 mm H) was placed around the set-up to prevent disturbance to the mantis.

A Kinobo USB B3 HD Webcam (Point Set Digital Ltd, Edinburgh, Scotland) was placed directly beneath the mantis to record behaviour. The output of the camera was fed to a Viglen genie PC computer and used as a monitor for an observer to record mantis behaviours off-line. The camera was positioned so that the observer only had a view of the mantis and not of the computer screen to ensure the observer coded the mantis behaviour blind to the stimuli presented.

### Behaviour

All behaviours were scored blind to the stimulus presented to avoid reporting bias.

We defined

1. *saccade as a* small head movement relative to the body line, in one direction
2. *tracking as a* head movement relative to the body line in the direction of the target made up of a number of saccades,
3. *peering* as a leaning from side-to-side with the head maintaining a fixed orientation toward the screen,
4. *strike* as rapid extension forelegs made toward the screen,
5. *optomotor response* as a leaning of the whole body in a horizontal direction without a separate head saccade or strike.

If the mantis responded in the direction the target was traveling it was defined as a response.

### Behavioural responses to wide-field motion (WFM) background, versus small-field motion (SFM) targets

Wide field motion was created by drifting sinewave gratings (subtending 140° by 128° at the mantis eye) presented. Small-field motion was created by a small portion of the drifting grating visible through a window that moved at the same speed and direction as the drift of the gratings. The window was an elliptical Gabor patch (a sinusoidal grating within a Gaussian envelope(Lee, 1996)) that subtended 23° by 11.65° at the eye, when measured between 50% transparency points within the target’s smoothed edge. Each mantis saw 20 combinations of motion type (WFM, SFM, each with 9 pairs of different spatial frequency (Sf) and temporal frequency (Tf), plus two control stimuli consisting of uniform dark areas of the same size as the Gabor patch and moving at velocities of 40 or 160 ° s^-1^ (these are shown in Figure 1). Each of the 20 combinations were presented 30 times to 6 mantises giving a total of 600 presentations per mantis. For the wide-field motion, each combination of spatial and temporal frequency was displayed two times within a block of trials, once traveling left and once traveling right. Targets appeared randomly at either the left or right side of the screen and travelled at a constant speed across the screen. All targets were the same mean luminance and moved over a homogenous grey background. Each presentation lasted a total of 5 seconds so, for example the WFM for each presentation the sinusoidal grating moved either left or right for a total of 5 seconds. In between trials, the mantis was aligned with the centre of the screen, using an alignment stimulus to ensure each test condition passed through the foveal region of the eyes (Nityananda et al., 2015). The alignment stimulus was a dark circle moving across the background image in a spiral motion from the edge of the screen to the centre, which attracted the mantises’ gaze and ensured that the head was finally positioned towards the centre of the screen. An observer classified each of the videoed reactions of the mantis without knowledge of the visual stimulus delivered.

**Figure 1.**
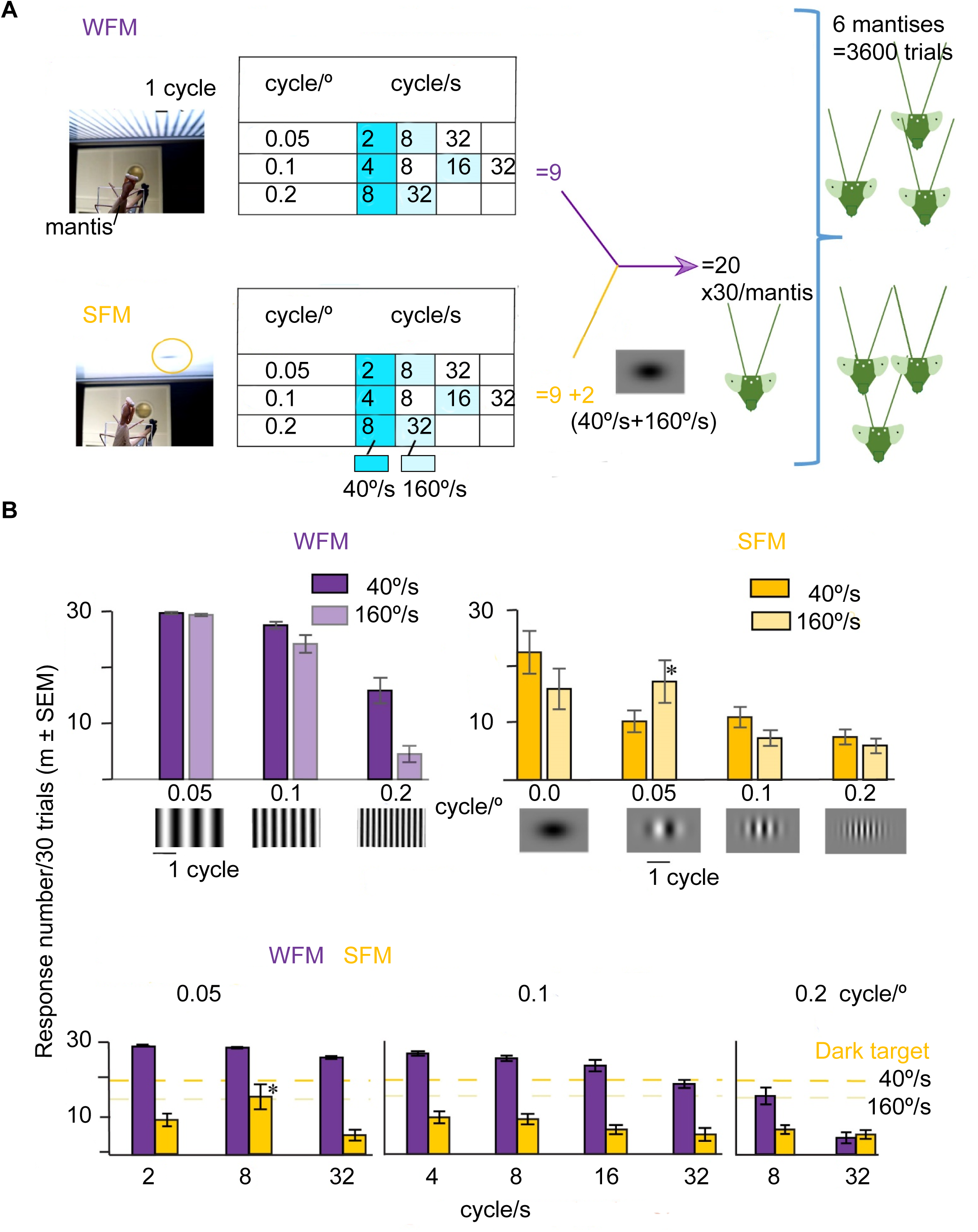
Behavioural responses of the mantis *Sphodromantis lineola* to wide-field motion (WFM), versus small-field motion (SFM) of sinusoidally modulated stimuli matched in their spatial and temporal frequencies. (A) Mantises responded with optomotor following responses to motion of a stripy sinewave grating (WFM top picture and Movie 1) or with tracking responses to motion of a small fuzzy Gabor window opened into the moving sinewave grating, the window moving with the grating (SFM bottom picture and Movie 2). To manipulate spatial and temporal frequencies independently, overall motion speed (measured in ° s^-1^) was made up by 9 different combinations of Sf (Spatial frequency in stripe cycles °^-1^), or Tf (Temporal frequency measured as cycles s^-1^). Both WFM and SFM had these same 9 combinations, as shown in the table for each motion type. SFM had two additional control trials with a solid dark target moving at either 40 ° s^-1^ s or 160 ° s^-1^, giving a total of 11 conditions for SFM. Three of the 9 spatio-temporal pairings had a speed of 40 ° s^-1^ (aqua squares), and three had a speed of 160 ° s^-1^ (light aqua squares). Values on the same horizontal line in the table had the same Tf and those on the same vertical line had the same Sf. Responses shown are mean response numbers given to each stimulus by the 6 mantises. For tracking, a response is counted if the mantis correctly tracked 1-4 of the four target transits across the screen, constituting the presentation (Movie 2). (B) Effect of Sf on responses to motion speed depends on the type of stimulus, **Left panel**, Optomotor responses to WFM stimuli. Optomotor responses depend on pattern Sf at both image speeds tested: 40 ° s^-1^ (GEE, 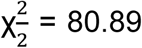, P< 0.001) and 160 ° s^-1^ (GEE, 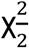, P< 0.001). **Right panel**, Tracking responses to SFM targets did not depend on pattern Sf but on the target type (a solid dark target as in the first two bars, Sf =0), or a sinusoidally modulated target GEE, 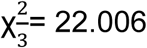, P< 0.001), with the solid dark target more likely to be tracked (Difference, *post hoc*, P= 0.003). One modulated target (asterisk) was tracked well, but this was a target with a Sf 0.05 cycles°^-1^, which results in 1.15 cycles over the 23 degree target, essentially a single light-dark object. (C) Comparison of the number of optomotor responses (purple bars) to WFM, and tracking responses (yellow bars) to SFM stimuli when stimuli were matched in Sf (X axis) and Tf (Y axis). We used 3 different Sf: 0.05 **(left panel)** 0.1 **(middle panel)** or 0.2 **(right panel)** cycles °^-1^ and a range of Tf from 2-32 cycles s^-1^. For reference the dark orange dashed-line on each graph indicates the mean number of trials the mantises tracked a dark unmodulated target moving at 40 ° s^-1^ and the light orange dashed line a dark unmodulated target moving at 160 ° s^-1^. Mantises tracked dark unmodulated SFM targets better than all but one SFM modulated target (asterisk). This was a target with a Tf of 8 cycles s^-1^, and a Sf 0.05 cycles °^-1^, giving a speed of 160 ° s^-1^. This Sf results in 1.15 cycles over the 23 degree target, essentially a single light-dark object. The mantis was more likely to display the optomotor response to the WFM of the modulated stimuli compared to a tracking response to a comparable modulated SFM stimulus (GEE, 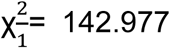, P< 0.001). The optomotor system (purple bars) is more easily triggered by drifting gratings than the tracking system (yellow bars) is by the small moving Gabor patches. The statistical relationship of the optomotor and tracking responses to Sf and, or Tf of both motion types was modelled with a GLM and results reported in the text and in Table 1.

We recorded the mantis’s behaviour as tracked (left or right) or optomotor response (left or right). Once a mantis’s response was recorded, the alignment stimulus was presented to align the mantis’s head ready for the next test stimulus. The stimulus type (Gabor patch or wide-field sinusoidal grating), the direction the stimulus travelled, and behavioural observations for each trial were recorded for later analysis. All data are shown either as the mean (± s.e.m.) response number or as the mean (± s.e.m.) response proportion.

**Table 1.** GLM: Univariate Analysis of Variance of the Mantises (A) optomotor responses to WFM; (B) Correct tracking responses to sinusiodally modulated bug-shaped stripy SFM with comparable spatial and temporal frequency to the WFM (C) The number of individual saccades contributing to the correct tracking responses to sinusiodally modulated bug-shaped stripy SFM of comparable spatial and temporal frequency to the WFM.

| Source | Type III<br>Sum of<br>Squares | df | Mean<br>Square | F | Sig. |
| --- | --- | --- | --- | --- | --- |
| Corrected Model | 16953.4<sup>a</sup> | 18 | 941.9 | 227.6 | .000 |
| Intercept | 2482.7 | 1 | 2482.7 | 599.9 | .000 |
| Animal(1-6) | .0 | 1 | .0 | .0 | .956 |
| Stimulus (SFM/WFM) | 11238.7 | 1 | 11238.7 | 2715.7 | .000 |
| Spatial Freq (SF) | 1005.4 | 2 | 502.7 | 121.5 | .000 |
| Temporal Freq (TF) | 272.8 | 4 | 68.2 | 16.5 | .000 |
| Stimulus * SF | 1005.4 | 2 | 502.7 | 121.5 | .000 |
| Stimulus * TF | 272.8 | 4 | 68.2 | 16.5 | .000 |
| SF * TF | 56.3 | 2 | 28.2 | 6.8 | .002 |
| Stimulus * SF * TF | 56.3 | 2 | 28.2 | 6.8 | .002 |
| Error | 368.3 | 89 | 4.1 |  |  |
| Total | 31058.0 | 108 |  |  |  |
| Corrected Total | 17321.7 | 107 |  |  |  |

a. R Squared = .979 (Adjusted R Squared = .974)

| Source | Type III<br>Sum of<br>Squares | df | Mean<br>Square | F | Sig. |
| --- | --- | --- | --- | --- | --- |
| Corrected Model | 2383.4<sup>a</sup> | 18 | 132.4 | 14.0 | .000 |
| Intercept | 563.1 | 1 | 563.1 | 59.7 | .000 |
| Animal(1-6) | 36.7 | 1 | 36.7 | 3.9 | .052 |
| Stimulus (SFM/WFM) | 1507.0 | 1 | 1507.0 | 159.8 | .000 |
| Spatial Freq (SF) | 56.2 | 2 | 28.1 | 3.0 | .056 |
| Temporal Freq (TF) | 142.2 | 4 | 35.6 | 3.8 | .007 |
| Stimulus * SF | 58.5 | 2 | 29.3 | 3.1 | .050 |
| Stimulus * TF | 157.1 | 4 | 39.3 | 4.2 | .004 |
| SF * TF | 67.4 | 2 | 33.7 | 3.6 | .032 |
| Stimulus * SF * TF | 65.0 | 2 | 32.5 | 3.5 | .036 |
| Error | 839.5 | 89 | 9.4 |  |  |
| Total | 5073.0 | 108 |  |  |  |
| Corrected Total | 3222.9 | 107 |  |  |  |

a. R Squared = .740 (Adjusted R Squared = .687)

| Source | Type III<br>Sum of<br>Squares | df | Mean<br>Square | F | Sig. |
| --- | --- | --- | --- | --- | --- |
| Corrected Model | 9414.1<sup>a</sup> | 18 | 523.0 | 5.8 | .000 |
| Intercept | 1735.7 | 1 | 1735.7 | 19.3 | .000 |
| Animal (1-6) | 96.7 | 1 | 96.7 | 1.1 | .302 |
| Stimulus (SFM/WFM) | 4891.7 | 1 | 4891.7 | 54.5 | .000 |
| Spatial Freq (SF) | 280.8 | 2 | 140.4 | 1.6 | .215 |
| Temporal Freq (TF) | 958.3 | 4 | 239.6 | 2.7 | .037 |
| Stimulus * SF | 282.1 | 2 | 141.1 | 1.6 | .213 |
| Stimulus * TF | 989.8 | 4 | 247.5 | 2.8 | .033 |
| SF * TF | 425.4 | 2 | 212.7 | 2.4 | .099 |
| Stimulus * SF * TF | 417.4 | 2 | 208.7 | 2.3 | .104 |
| Error | 7986.8 | 89 | 89.7 |  |  |
| Total | 23521.0 | 108 |  |  |  |
| Corrected Total | 17400.9 | 107 |  |  |  |

a. R Squared = .541 (Adjusted R Squared = .448)

### Increasing the number and closeness of internal features decreased the number of trials in which a chequered target was tracked

Targets were small 80 × 80 pixel squares subtending 19.3 × 19.3° at the eye and were either homogenously dark (0.052cd m^-2^), or a chequerboard whose internal square chequers were 5, 10 or 20 pixels (1.2, 2.4 and 4.8 degrees) in width. In all cases the chequerboard targets moved over a grey background that matched the target’s mean luminance. Targets had equal numbers of white (cd m^-2^) and dark (0.052 cd m^-2^) chequers. In each trial, the target appeared randomly at the left or right side of the screen, and then travelled across the screen horizontally before returning; this cycle of motion was repeated four times for 10 seconds. The target moved with a sinusoidal function with a maximum speed of 1167 pixels s^-1^ (209°s^-1^) reached when the target was at 0, in the centre of the screen. When the target position reached -1 or 1 (either edge of the screen) velocity was zero. Targets reversed direction off the screen. Motion of the square was slower at the sides of the monitor and faster in the middle to ensure target speed over the eye was constant. Each target type was displayed 15 times to each mantis, with a sample size of 10 mantises.

### Visual stimuli developed to test if target tracking is affected by motion of the background

Backgrounds of random chequer-board of three different chequer sizes were tested. The target, if present, was a dark 80 × 80 pixel square (0.052 cd m^-2^) which moved across a central grey strip, 80 or 329 pixels in height. 10 pixels subtend 2.4° on the eye. All background patterns had the same mean luminance (36cd m ^-2^), and had equal numbers of white (72 cd m^-2^) and black chequers (0.052 cd m^-2^). The uniform grey strip matched this mean luminance. The position and speed of the target varied sinusoidally and was at its maximum (209 °s^-1^ or 1167 pixels s^-1^), when it was at 0 (the centre of the screen). The target appeared randomly at the left or right side of the screen, and then travelled back and forth across the screen a total of four times. Tracking was scored if one of these 4 movements were correctly tracked. The target was not visible to the mantis at either edge of the screen when it changed direction. All data are shown as the mean (± s.e.m.) response number or response proportion.

### Quantification and Statistical Analysis

Statistical analysis was carried out using several methods available in SPSS v 22-23, the method depending on the complexity of the data to be analysed or modelled. The level for significance was set at p <0.05. For simple analysis we used generalized estimating equations (GEE, binary logistic model) in SPSSv22. With background movement type (still, in-phase and out-of-phase) and background pattern (5-, 10- and 20-pixel) as the independent variables. The number of trials where tracking occurred (out of the total of 15 presentations for each condition) was used as the dependent variable. Mantis was a repeated factor. For more complex analysis with interactions between variables we used Generalised linear models (GLM) in SPSS v23. Unless stated, animal was included in the model as a random term to check effects were the same across all individual mantises, particularly as new individuals were added over the time course of the experiment. We checked the assumptions for the model to be correct: predicted values of the residuals were clustered with no systematic trend in variance and the residuals were normally distributed (Everitt and Dunn, 2001). Intercept was included in the model.

Statistical analysis was carried out using several methods available in SPSS v 22-23, the method depending on the complexity of the data to be analysed or modelled. The level for significance was set at p <0.05. For simple analysis we used generalized estimating equations (GEE, binary logistic model) in SPSSv22. With background movement type (still, in-phase and out-of-phase) and background pattern (5-, 10- and 20-pixel) as the independent variables. The number of trials where tracking occurred (out of the total of 15 presentations for each condition) was used as the dependent variable. Mantis was a repeated factor. For more complex analysis with interactions between variables we used Generalised linear models (GLM) in SPSS v23. Model 1 In the model, optomotor and tracking responses were included as dependent variables; and animal (1-6), pattern spatial frequency (SF), and temporal frequency (TF) were included as fixed factors. The intercept was included in the model. Model 2 Unless stated, animal was included in the model as a random term to check effects were the same across all individual mantises, particularly as new individuals were added over the time course of the experiment. We checked the assumptions for the model to be correct: predicted values of the residuals were clustered with no systematic trend in variance and the residuals were normally distributed (Everitt and Dunn, 2001). Intercept was included in the model.

## Results

### Comparison of Responses to Wide-Field and to Small-Field Motion Stimuli

Moving striped patterns such as sinusoidally-modulated luminance-gratings are a good way of studying the sensitivity of the visual system to the luminance changes accompanying image motion, because their luminance at the eye changes regularly in time and in space. Optomotor responses that stabilize the animal relative to its surroundings are generated by correlation based motion-detectors in the visual system (Reichardt and Guo, 1986), their hallmark is that with regular repeating stripy patterns (Fig. 1A, B) such as a sinusoidally-modulated luminance-grating their speed tuning is determined by a combination of the pattern’s temporal frequency, and spatial frequency according to the ratio

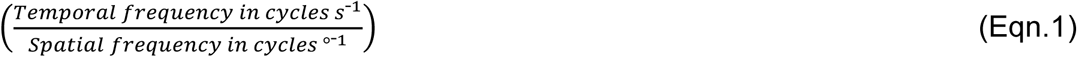

(Kulikowski and Tolhurst, 1973; O’Carroll et al., 1997; Reichardt and Poggio, 1979; Thompson, 1982). Sinusoidally modulated luminance gratings could also reveal if classical, correlation based motion-detectors underlie small-target movement-detection, if the grating was small enough. Such target motion was created by making only a small portion of a larger moving sinewave grating visible through a window that moved at the same speed and direction as the drift of the grating behind it. The target, a Gabor patch looked like a fuzzy, stripy bug moving across the screen (Fig. 1). This target allowed us to compare tracking and optomotor responses to the same range of spatial and temporal frequencies of motion, to determine if the two responses depended in the same way on the ratio of temporal to spatial frequency. We included a small dark target with no stripes so we could compare the response to this, with the response strength to our stripy bugs (Fig. 1). In these experiments the mantis hung freely under a small Perspex platform and viewed a computer screen. In response to a wide-field movement over the whole monitor screen (Fig. 1), the mantis showed optomotor responses in which it followed the stimulus in a continual, smooth movement, generated by various body parts including leaning by the legs (Video S1) (Nityananda et al., 2015; Reichardt and Guo, 1986). In contrast, a typical tracking response consisted of head rotations, following movement of the small Gabor-target (Video S2) (Prete and Mahaffey, 1993; Prete and McLean, 1996). Mantises responded with optomotor following responses to motion of a stripy sinewave grating (WFM top picture and Video S1) or with tracking responses to motion of the small fuzzy Gabor window opened into the moving sinewave grating. To manipulate spatial and temporal frequencies of WF and SF motion independently, overall pattern motion speed (measured in ° s^-1^) was made up by 9 different combinations of Sf (Spatial frequency in stripe cycles °^-1^), and Tf (Temporal frequency measured as cycles s^-1^) these included three combinations giving motion speeds of 40 ° s^-1^ (aqua boxes in the table, Figure 1A) and three giving motion speeds of 160 ° s^-1^ (light-aqua boxes in the table, Figure 1A). Both WFM and SFM had these same 9 combinations. SFM had two additional control trials with a solid dark target moving at either 40 ° s^-1^ or 160 ° s^-1^ (aqua text), giving a total of 11 conditions for SFM. The six mantises saw 30 repetitions of each motion and the responses recorded. The results were analysed statistically by fitting a GLM but to get a feel for what they looked like they have been plotted as either, mean responses to the WF or SF stimuli plotted against motion spatial frequency (WFM: purple bars; SFM yellow bars, Fig. 1B), or as responses to both WF and SF stimuli, plotted against temporal frequency at each spatial frequency tested (WFM: purple bars; SFM yellow bars, Fig. 1C). The first thing to see is that a comparison of the number of optomotor responses (purple bars, Fig. 1) to WFM, and tracking responses (yellow bars, Fig. 1) to SFM stimuli showed the optomotor system is more easily triggered by drifting gratings than the tracking system is by the small moving Gabor patches. Also responses to WF and SF motion don’t both follow the same trends, as predicted, optomotor responses show a systematic relationship with Sf of the WFM in Figure 1B, declining as the spatial frequency increases at both speeds (40 and 160° s^-1^). Whereas in the same figure (yellow bars, Fig. 1B) tracking responses do not show a systematic shift as spatial frequency increases. One modulated target (asterisk) was tracked well, but this was a target with a Sf 0.05 cycles °^-1^, which results in 1.15 cycles over the 23 degree target, essentially a single light-dark object. Over the range of temporal frequencies we tested (Fig. 1C), both optomotor responses to WFM (purple bars) and tracking responses to SFM (yellow bars) decrease as Tf increases. For reference the dark orange dashed-line on the graphs indicates the mean number of trials the mantises tracked a dark unmodulated target moving at 40 ° s^-1^ and the light orange dashed line a dark unmodulated target moving at 160 ° s^-1^. The mantis can clearly track at these motion speeds. Mantises tracked dark unmodulated SFM targets better than all but one SFM modulated target (asterisk). This was a target with a Tf of 8 cycles s^-1^, and a Sf 0.05 cycles °^-1^, giving a speed of 160 ° s^-1^. This Sf results in 1.15 cycles over the 23 degree target, essentially a single light-dark object.

In Figure 2 the optomotor and tracking responses are each plotted against the ratio of

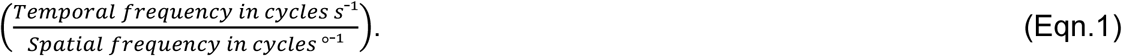

**Figure 2.**
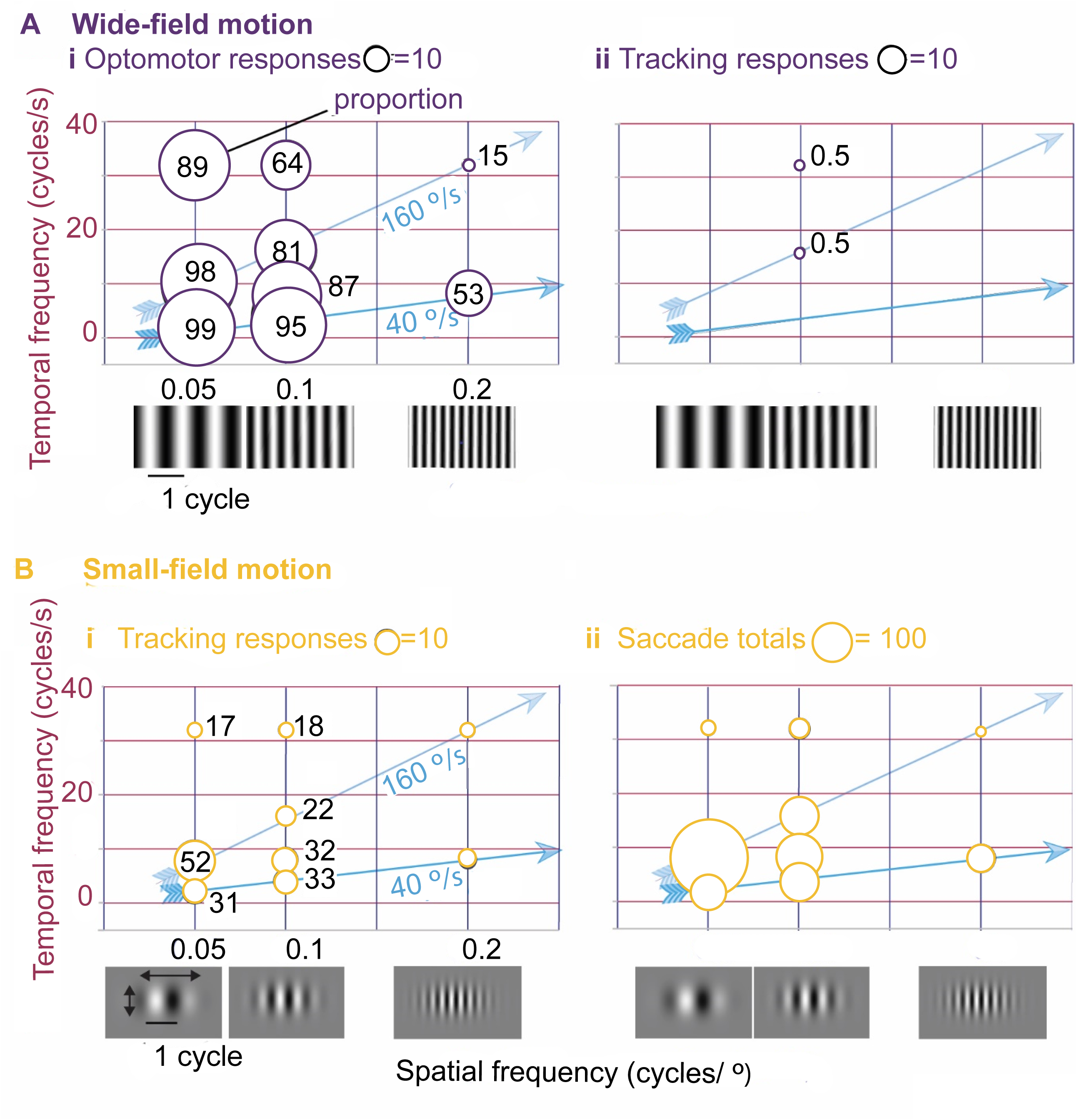
Optomotor and target tracking responses depend differently on a pattern’s spatial frequency and temporal frequency to judge motion speed. The data from the experiment described in Figure 1 has been replotted to show the overall mean response rates to the WFM compared with SFM stimuli when motion speed was made up by different combinations of Tf and Sf. Each combination of spatio-temporal frequency was displayed to 6 mantises a total of 30 times. The diameter of each circle represents mean number of optomotor or tracking responses given by the six mantises to these 30 presentations. The proportion of responses this represents is stated on the graph, inside each circle. Diagonal arrows show the responses to stimuli moving at a speed of 40 ° s^-1^, (dark aqua) or 160 ° s^-1^, (light aqua). (A) Optomotor and Tracking responses to WFM. Only two tracking responses were produced to 1620 WFM stimuli. (B) Tracking responses given and total transits tracked for all SFM targets. Each of the 30 presentations had 4 tracking opportunities as the target moved four times across the screen not all mantises followed all 4 screen crossings (transits). If any of the 4 transits was tracked the presentation was counted as tracked.

Plotting them this way allows us to see the optomotor and tracking responses of the same mantis, under the same conditions and to the same pattern speed. Pattern speed in ° s^-1^ is aligned along diagonal iso-speed lines (Fig. 2A and B): Dark aqua arrows connect responses to motion of 40 ° s^-1^ whereas light aqua arrows connect responses to WF and SF motion of 160 ° s^-1^. The mantises are sensitive to a broad range of WFM velocities, and were most sensitive, showing most optomotor responses when the spatial frequency was at the lowest of the range tested, around 0.05 cycles s^-1^, and the temporal frequency was below 8 Hz. The strength of the optomotor response shows a gradual decline when spatial or temporal frequencies increase, as has been shown previously. The greatest response was to a WFM speed of 40 ° s^-1^. In a recent study comparing human and mantis motion detection *(Nityananda et al., 2017; Nityananda et al., 2015)* mantises were most sensitive to relatively low spatial frequencies (between 0.01 and 0.1 cycles °^-1^). To our SFM the same mantises were again most sensitive to low spatial frequencies (between 0.05 and 0.1 cycles °^-1^) but the preferred temporal frequency was 8 cycles s^-1^. The greatest responses lay on the 160 ° s^-1^ speed line (Fig. 2B). The tuning was narrower than with the WFM, particularly obvious when the number of tracked transits of each presentation is taken into account by plotting how many of the 4 SFM transits were tracked rather than merely indicating that a correct tracking response was given to one of the four transits (right graph, Fig. 2B). The responses are bunched at 160 ° s^-1^, which corresponds to a stimulus temporal frequency of 8 cycles s^-1^ and spatial frequency of .05 cycles °^-1^, essentially a single dark stripe making up the moving bug. Suggesting a suppressive lateral interaction occurs between stripes in the motion-detectors underlying target tracking.

The mantises’ responses to WF and SF motion, were compared statistically by fitting responses using a generalized linear model (GLM, Table 1). Optomotor and tracking responses to WFM and SFM respectively, were the dependent variables; pattern spatial frequency (Sf), temporal frequency (Tf) and animal were included as fixed factors, and intercept was included in the model. For WFM, both spatial frequency and temporal frequency of the pattern were significant factors in determining the mantises’ optomotor responses (Table 1A, Spatial frequency: Type III Sum of Squares (SS), 1005, df 2, Mean Square (MS) 503, Sig.0.000; Temporal frequency: SS 256.85, df 4, MS 256.85, F 68, Sig. 0.000). However, to SF Gabor-target motion, only the temporal frequency of the target was significant in determining tracking responses, (Table 1B, Spatial frequency SS, 56, df 2, MS 28, Sig.0.056; Temporal frequency: SS 96.31, df 4 MS 36, F 4, Sig. 0.007) and the number of saccades in the tracking responses (Table 1C, Spatial frequency: SS, 280.8, df 2, Mean Square (MS) 140.4, Sig.0.215; Temporal frequency: SS 958.3, df 4, MS 239.6, Sig. 0.037).

For both optomotor and tracking responses, there was a significant interaction between spatial and temporal frequency (Table 1A-C). Animal was not a significant factor in either optomotor or tracking responses. The spatial frequency of both wide-field and Gabor targets were well above the spatial resolution of a single ommatidium. In the praying mantis inter-ommatidial and photoreceptor acceptance angles depend on the state of light adaptation and the time of day; for example photoreceptors have mean acceptance angles of 0.74° (S.D. = 0.1) when light-adapted, and 1.1° (S.D. = 0.2) when dark-adapted in daytime (Rossel, 1979). This angle sets the limit to spatial resolution and is much finer than the 5, 10 or 20° per light/dark cycle used here, which means that the lack of tuning to spatial frequency is not caused by limits at the retina.

### Evidence for short-range inhibition: addition of internal features to a target decreases tracking

We tested whether increasing the number and closeness of a target’s internal light and dark features affected the number of small-field motion (SFM) targets tracked. In these experiments SFM targets were small squares subtending 14.32° x 14.32° at the eye (80 × 80 pixels) and were either homogenously dark (0.052cd m^-2^), or a chequerboard whose internal square chequers were 1.2, 2.4 and 4.8° (5, 10 or 20 pixels) wide (Fig. 3). In all cases the chequerboard targets moved over a grey background that matched the chequered target’s mean luminance. Chequered targets had equal numbers of white (72 cd m^-2^) and dark (0.052 cd m^-2^) chequers arranged randomly. Responses to these stimuli were not shaped by the limit of spatial or temporal resolution of the mantis. Even the finest, 5 pixel, 1.2° sized chequer-board squares were above the 0.7° resolution of a single foveal ommatidium (Rossel, 1979) and at an angular size responded to by target-selective selective neurons of the in the lobula complex of the mantis *Tenodera aridifolia* (Yamawaki, 2019; Yamawaki and Toh, 2003) as shown in Figure 3 A of Yamawaki and Toh (2003). In our experiments motion of the target had a maximum speed of 209° s^-1^, which is within 20% of the highest contrast sensitivity of the mantis optomotor response (Nityananda et al., 2015) and close to the angular velocity giving maximum response, of target-selective neurons of the mantis *Tenodera aridifolia* to dark moving squares (Yamawaki and Toh, 2003) as in Figure 3 B in Yamawaki and Toh (2003).

**Figure 3.**
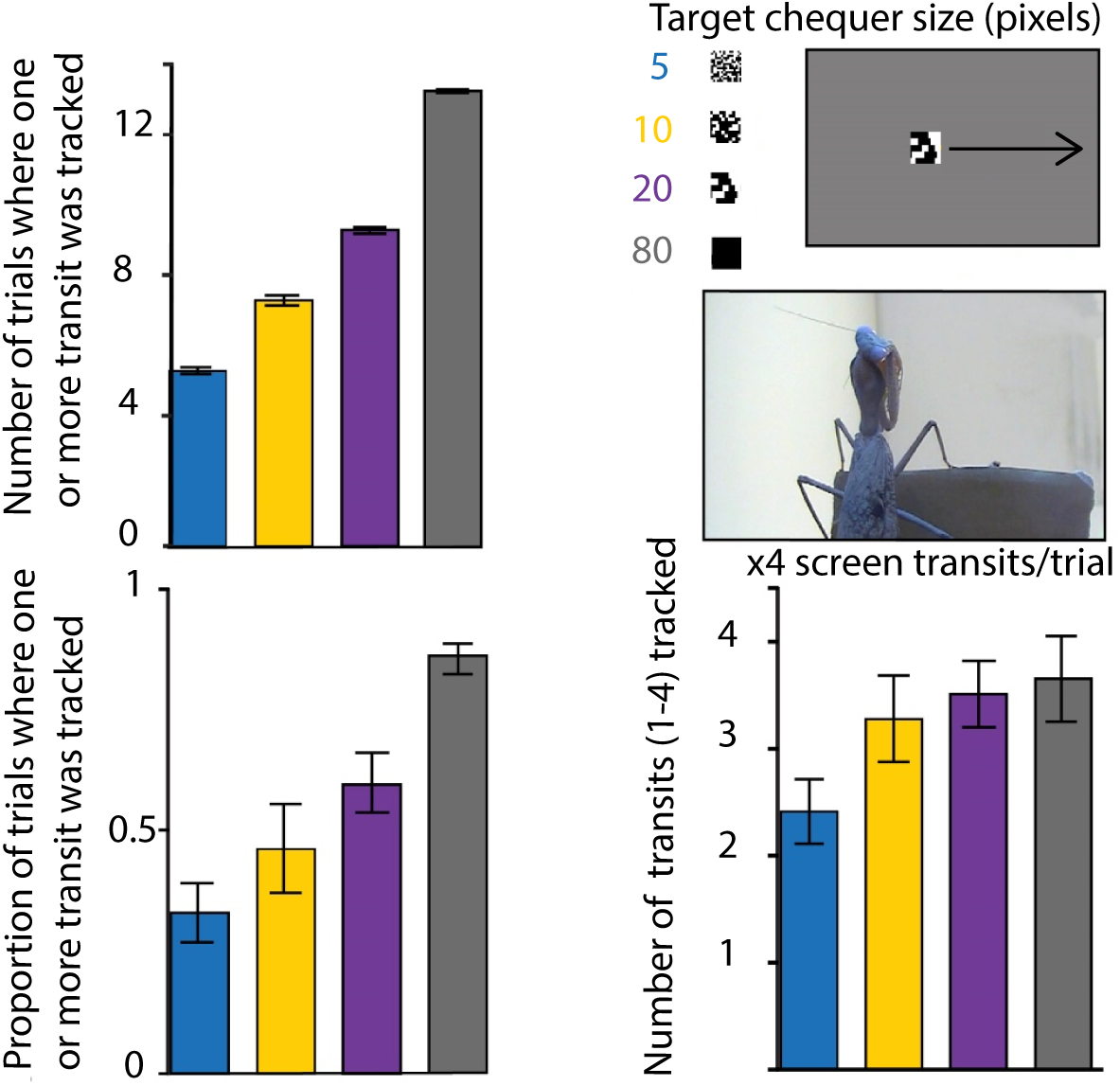
Increasing the number and closeness of target internal features decreased the number of trials the mantis tracked. (A) Number of the trials in which the target was tracked during the 4 cycles of target motion. (B) Proportion of the trials in which the target was tracked. (C) Number of individual cycles tracked. Visual targets were created with a custom written script using Matlab (MathWorks) and the Psychophysics Toolbox. The test stimulus consisted of a static grey background image and a square target (80 pixel X 80 pixel) subtending 19.3°on the mantis retina. In each trial, the target appeared randomly at the left or right side of the screen, and then travelled across the screen horizontally before returning; this cycle of motion was repeated four times for 10 seconds. The target moved with a sinusoidal function with maximum speed of 1166.7 pixels s^-1^ at the centre of the screen. The target was not visible to the mantis at either edge of the screen when it changed direction. This ensured from the mantids perspective that the target appeared to move at a constant speed across the screen. For further stimulus details see Methods section.

The probability that the mantises tracked targets was negatively affected by the target internal pattern (dark, 20, 10, 5 pixel random chequers; Fig. 3) (General Estimating Equation, GEE, 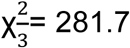, P<0.001; Fig. 2). Mantises were more likely to track a completely dark target over patterned targets (Helmertz *Post hoc, P*<0.001; Fig. 2). Comparing the patterned targets, mantises were more likely to track a 20 pixel patterned target compared to the 5 pixel and 10 pixel patterned target (GEE, pairwise *Post hoc, P*<0.001, P<0.001; Fig. 3). Mantises struck at these targets but strike numbers were low to all stimuli, <5 per 150 stimuli, because the targets were 60 mm away, out of normal physical catch range (Nityananda et al., 2018). If classical correlation based motion-detectors were providing input to the tracking response, we would expect the addition of more edges to a small target, once the resolution of the single ommatidium (0.7°) is exceeded (Rossel, 1979), would increase target tracking but this was not what we found. On the other hand, with non-classical motion-detectors, adding fine detail to the target, particularly closely spaced light and dark edges reduces the detector’s response even before the physical limit to spatial resolution or response size is reached (Nicholas et al., 2018; Simmons and Rind, 1992; Simmons and Rind, 1997). This was what we had already observed in the tracking responses to Gabor-targets in Figure 1B, where adding detail to the pattern, by increasing spatial frequency, reduced the tracking response, except when, as explained in the text, with a spatial frequency of 0.05 cycles °^-1^ the Gabor target was essentially a single dark stripe.

### Evidence for long-range inhibition: patterned WF background supresses object tracking

To investigate evidence of longer-range spatial antagonism or lateral inhibition, a feature of non-classical motion detection such as locust looming detectors (Rind and Simmons, 1998), we presented different wide-field backgrounds, and targets together in combinations with and without background motion (Fig. 4 A, Bi), or, with and without targets (Fig. 4 A, Bii-iii). In the first condition, which acted as a control, tracking responses of the mantises were recorded to motion of a dark target over a uniform grey background (grey line, Fig. 4C). This response could then be compared to responses when the same target motion occurred over a grey strip between two stationary randomly chequered backgrounds of the same mean luminance as the grey strip (Fig. 4 Bi). In this situation the mantises own tracking motion would lead to motion of the background over its eyes. We used backgrounds of 5, 10 and 10 pixel chequer sizes, to see what effect that would have on any response suppression.

**Figure 4.**
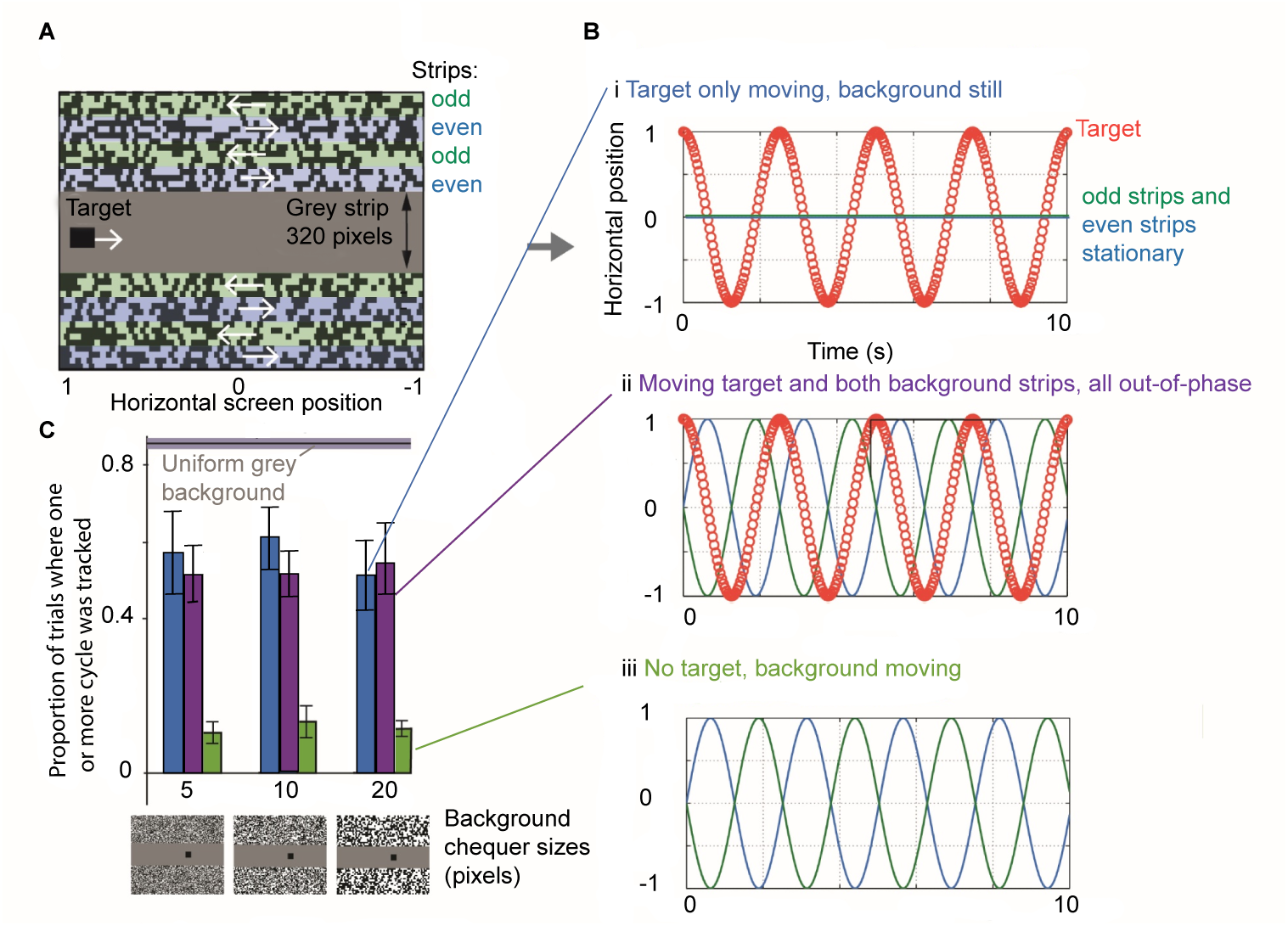
Target tracking is reduced by motion of the background. (A) To produce background motion without causing an optomotor response that could interfere with target tracking, motion of the background was split into eight numbered strips, each 20 pixels in height. Odd and even strips of the random chequerboard background moved in opposite directions, 180° out-of-phase, (white arrows). (B) Experimental conditions: target motion (white arrow) was (i) against a stationary background, (ii) out-of-phase to background motion. Or, (iii) there was no target motion just background motion as in (ii). (C) The proportion of the trials tracked in i-iii, is plotted as a bar chart and is compared to the response given to a target moving over a uniform grey background. 9 Mantises contributed data, 7 received 15 trials of each stimulus in a random order, one completed 10 trials, and one completed 5 trials. The mean (+/- s.e.m) proportion of times the mantids displayed a tracking response to each condition is shown. Each combination of visual condition and pattern element size were displayed 15 times to each mantis.

We next tested the effect of a distant moving background on tracking. We designed motion of the background chequerboard patterns to ensure no overt optomotor behaviour was caused, and which still allowed the effect of background motion on tracking to be determined. To do this each chequerboard of the background consisted of alternating strips, each 20 pixels in height (referred to as ‘odd’ and ‘even’ strips). All even strips moved in phase with each other, and all odd strips moved in phase, but odd and even strips moved 180°out-of-phase with one another (illustrated in Fig. 4A, Bii). The random placement of chequers in the strips can lead to aggregations of dark squares which look like a dark object. To control for this effect, we measured the tracking response to motion of the background with no central dark target present (Fig. 4, Biii; and green bars in C). Mantises a showed low proportion of tracking responses across all background chequer sizes 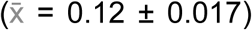 even in the absence of a moving central dark target (Fig. 4Biii and green bars C). However these tracking responses with no central target were significantly reduced compared to those where a central target moved over either a still background (Fig. 4Bii and blue bars C) (GEE, pairwise post hoc, P<0.001) or a moving background (Fig. 4Bi and magenta bars Fig. 4C) (GEE, pairwise post hoc, P<0.0001). The number of trials tracked was significantly different under the three relative motion conditions i-iii (GEE, 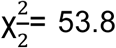, P<0.001). That the tracking response could be reduced when WFM was self-generated shows that the inhibition is produced regardless of whether the WFM is generated by the mantis itself or is generated externally (Fig. 4Bii and blue bars C) and does not require there to be overt optomotor behaviour. Suggesting it is a feature of the target tracking process itself.

### Inhibitory interactions: patterned WF background supresses object tracking in proportion to their physical separation

Wide field, moving gratings inhibit responses by the LGMD1and LGMD2 neurons both to small, translating objects (Rowell et al. 1977; Pinter 1977, 1979) and to approaching objects (Rind and Simmons 1992; Simmons and Rind 1997). The suppression of LGMD1 and 2 responses depended on the spatial features of the WFM, showing a peak suppression with drifting gratings of 20° in period, moving at a velocity between 100 and 200° s^-1^. In those experiments the locust was fixed in position. To quantify any such long range inhibitory interactions in the mantis we used three motion conditions (Fig. 5A): i, target motion with a stationary background where background motion was self-generated; ii, target motion was 50% in phase and 50% out of phase with background motion and iii, target and background motion were out of phase. We also looked at spatial effects of the WFM on target tracking. The spatial features, (chequer sizes) of each background ranged between 5-20 pixels.

**Figure 5.**
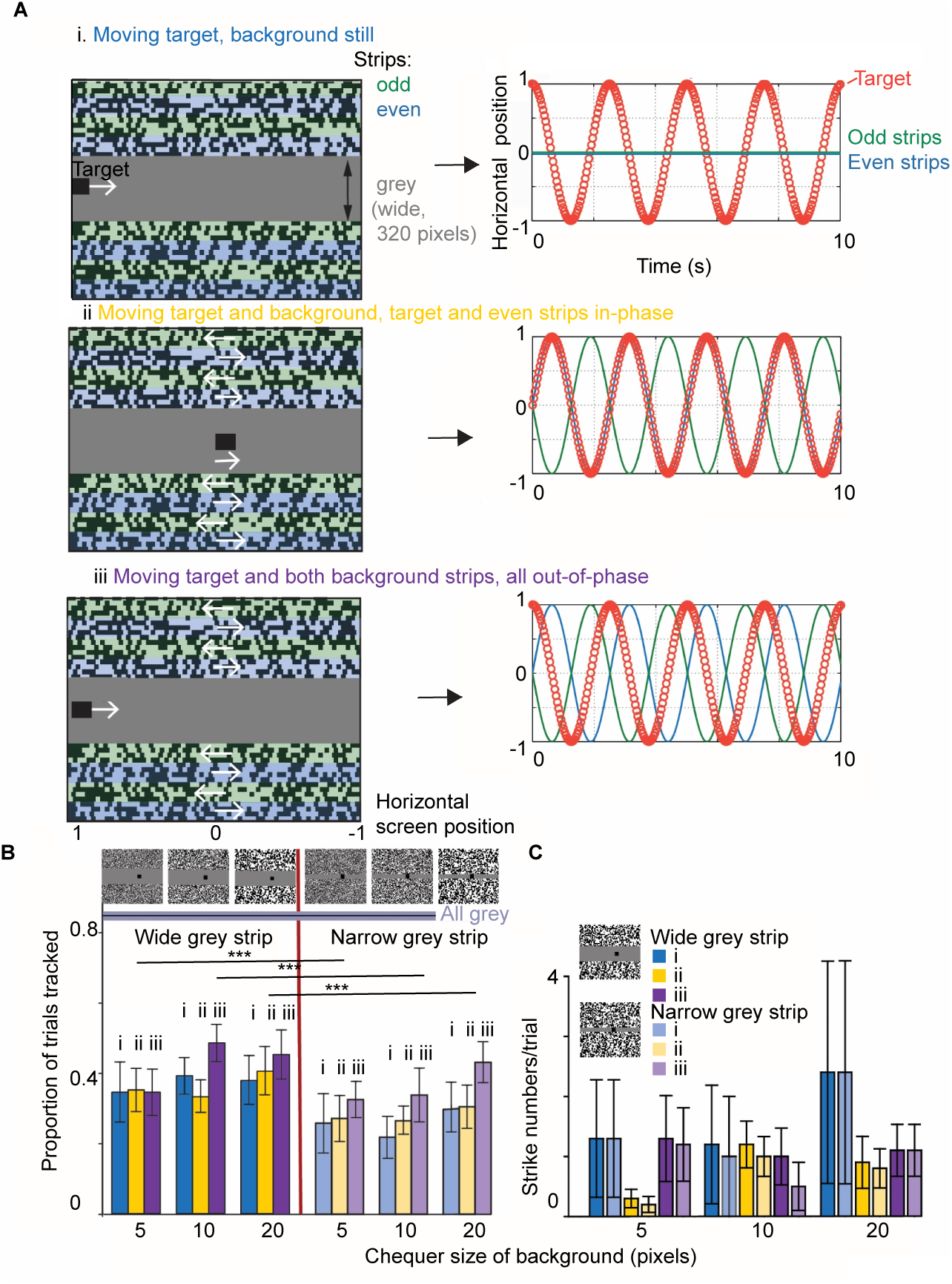
A patterned WF background supresses object tracking and striking in proportion to its proximity to the target, regardless of the phase of their relative motion. (A) Motion of the background was split into eight numbered strips, each 20 pixels in height. All odd or all even numbered strips moved in the same direction. Odd (green lines) and even (blue lines) strips moved 180° out-of-phase to one another. Three relative motions of target and background were presented to each mantis illustrated as it appears on the monitor (left panel), with white arrows indicating initial motion direction. (i), target motion with a stationary background, (ii), target and background motion 50% in-phase, matching the movement of the odd rows of the background, (iii), target and background motion out of-phase, not matching the movement of even or odd rows of the background. (B) Tracking responses to these three motion conditions. The central grey strip was either 320 pixels or 80 pixels in height. Increased proximity of the background (right side of the graph) significantly reduced the proportion of trials tracked. A horizontal line indicates the high proportion of trials tracked when targets moved over a uniform grey background. Motion of the background alone accounted for 0.12 ± 0.02 s.e.m. of the trials tracked, across all three sizes of chequers. (C) Strike numbers also reduced with increased proximity of the background to the target. Data from this experiment was fitted using a GLM (Table 2). Of the original 10 mantises used in (B), 5 mantises died during the experiment so these only saw one background separation so 5 additional mantises were recruited for the new tests with the narrow target background separation. Each mantis received 15 trials of each stimulus configuration. 9 out of the 10 mantises contributed to experiments on both chequered and grey backgrounds. Experimental details were as in Figure 4.

For all three motion conditions and background spatial features, we tested two physical separations of WF background and target, (Fig. 5B). WFM of any sort reduced the proportion of trials the mantises tracked: a dark moving target on a uniform grey background was tracked 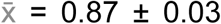, and addition of any random-chequered background led to a mean tracking proportion of 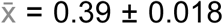, a 48% reduction across all conditions for a wide target background separation. And 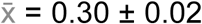, a 56% reduction across all conditions for a narrow target background separation. Increasing the proximity of a WFM stimulus significantly reduced the number of tracking responses and the proportion of targets tracked (Fig. 5B left compared to right panel). A GLM was fitted to the tracking data from this experiment (Table 2A). One factor was highly significant, the separation of WF background and target, such that as separation decreased, tracking was supressed (Fig. 5B). Two other factors were significant both alone and in their interaction: the WF background chequer size (Table 2B) and the relative motion condition of background and target (i-iii Table 2C). Although there was no significant difference between tracking responses to motion of the WF background in-phase with target motion and to motion of the target over a stationary background both background conditions supressed tracking significantly more than out-of-phase (i-iii Table 2C; and compare yellow and blue with magenta bars in Fig. 5C). Of course, as the mantis was free to move when it tracked the SF target near a background the mantises own tracking will also move the whole chequerboard background across its eye in the opposite direction to target motion with a speed generated by its tracking motion. Although this complicates the interpretation of the different phases of motion between target and, both the odd and even background strips, it does not change the increased suppression of target tracking observed as the distance between target and background decreased. An effect observed over a range of different WF motion conditions.

**Table 2.** GLM fitted to tracking data, that is the number of tracking responses to targets and chequered backgrounds as plotted in Figure 5B. The dependent variable was number of tracking responses to 15 stimulus presentations. The fixed factors were: 1. proximity of the WF background to the target (no separation of target and background, compared with 120 pixels separation of target and background); 2. Background motion condition (i, Still WF background, ii, WFM 50% in-phase with target motion and iii, WFM out-of-phase with target motion); and 3. WF background chequer size, (5, 10 or 20-pixels^2^). Mantis was added as a random factor to take account of variability between individuals. **(A)** fixed effects. Each of the fixed effects: proximity of the WF background to the target; background motion condition (i, Still WF background, ii, WFM 50% in-phase with target motion and iii, WFM out-of-phase with target motion); and WF background chequer size, each have a significant effect on trials tracked. There is an interaction between background condition and chequer size. **(B)** pairwise comparisons of trials tracked with different background chequer sizes (pixels^2^), **(C)** pairwise comparisons between background condition and trials tracked (0, Still; 1, WFM 50% in-phase with target motion and 2, WFM out-of-phase with target motion). Results that are significant at the 0.05 level are highlighted in bold.

A Type III Tests of Fixed Effects
| Source | Numerator df | Denominator df | F | Sig. |
| --- | --- | --- | --- | --- |
| Intercept | 1 | 14.69 | 12.26 | .003 |
| Background proximity | 1 | 171.50 | 16.17 | **.000** |
| Background condition | 2 | 164.68 | 3.97 | **.021** |
| Background chequer size | 2 | 164.68 | 4.14 | **.018** |
| Background condition *<br>Background chequer size | 4 | 164.68 | 3.02 | **.019** |

B Pairwise comparisons between background chequer sizes (5, 10 and 20 pixel),
| (I) Background<br>chequer size<br>(pixel<sup>2</sup>) | (J) Background<br>chequer size (pixel<sup>2</sup>) | Mean<br>Difference (I-J) | Std.<br>Error | Df | Sig. |
| --- | --- | --- | --- | --- | --- |
| 5 | 10 | -1.62 | 1.22 | 120 | .189 |
|  | 20 | **-4.50** | 1.22 | 120 | **.000** |
| 10 | 5 | 1.62 | 1.22 | 120 | .189 |
|  | 20 | **-2.88** | 1.22 | 120 | **.020** |
| 20 | 5 | **4.50** | 1.22 | 120 | **.000** |
|  | 10 | **2.88** | 1.22 | 120 | **.020** |

| (I) Background condition<br>(i=stationary,<br>ii=50% in phase,<br>iii=out-of-phase) | (J) Background condition<br>(i=stationary,<br>ii=50% in phase,<br>iii= out-of-phase) | Mean Difference (I-J) | Std. Error | Df | Sig. |
| --- | --- | --- | --- | --- | --- |
| i | ii | 1.23 | 1.22 | 120 | .315 |
|  | iii | **-2.75** | 1.22 | 120 | **.026** |
| ii | i | -1.23 | 1.22 | 120 | .315 |
|  | iii | **-3.98** | 1.22 | 120 | **.001** |
| iii | i | **2.75** | 1.22 | 120 | **.026** |
|  | ii | **3.98** | 1.22 | 120 | **.001** |

The increasing proximity of a WF stimulus to the target (dark compared with light bars in Fig. 5C) also reduced the overall number of strikes made at the targets after they were tracked, but there was a lot of individual variability in strike rates between the mantises.

## Discussion

### Tracking behaviour in the mantis involves a looming-type pathway

Our results argue that tracking behaviour in the mantis involves a looming-type pathway, most thoroughly studied in the locust, although also present in the mantis (Yamawaki and Toh, 2003; Yamawaki and Toh, 2009). Three features of a looming-type motion-detector were shown by target tracking behaviour in a mantis, first tracking responses did not depend on classical motion detection pathways involving elementary motion-detectors that compare inputs shifted in time and space to generate their directional responses. Tracking depended more on temporal aspects of motion rather than spatial input features (Table 1). Second, adding more detail inside a chequered target while maintaining its mean luminance, supressed tracking and lastly, simultaneous wide-field motion in any direction supressed tracking in proportion to the background’s physical distance from the moving target and in proportion to the detail of the background (Table 2). In looming-detectors, it is known that adding fine detail to an object, particularly closely spaced light and dark edges reduces output, as does motion of a nearby high contrast WFM stimulus (Nicholas et al., 2018; Simmons and Rind, 1992; Simmons and Rind, 1997). The suppression of locust LGMD1 and 2 responses, for example, depend on the spatial features of the WFM, showing a peak suppression with drifting gratings of 20° in period, moving at a velocity between 100 and 800°s^-1^. In our mantis experiments we observed maximum tracking suppression with the smallest WFM background chequer size: 5 pixels subtending 1.2° at the eye, all backgrounds moved at a high velocity, 1166 pixels s^-1^ (209° s^-1^) within the range of peak suppression in the locust (Simmons and Rind, 1997). Our backgrounds were high contrast and the random distribution of dark chequers could be aggregated together making a direct comparison to the sinewave WFM backgrounds used in the locust difficult, except that in both cases background motion speed was high and above 200° s^-1^.

### Both long and short-range suppression could derive from the same process in the looming-type motion detection

Maximum suppression of target tracking occurred when the size of the targets’ internal chequers was the smallest, suppression was significantly greater with chequers of 5 or 10 pixels compared with 20 pixels (Fig. 2). The long range lateral suppression of target tracking by a chequered background showed the same correlation with the pixel size of the backgrounds’ chequers, the greatest suppression occurring with the 5 pixel chequers (Fig. 5B and Table 2). Suggesting both types of suppression could derive from the same process in the looming-type motion detection as has been suggested for looming detection by the LGMD 1 and 2 neurons in the locust (Rind and Simmons 1997; Rind *et al* 2016; Cocks PhD thesis, 2020). Connectomics studies of the retinotopic TmA inputs to the LGMD 1 and 2 show widespread connectivity between afferents from several retinal columns, with particular hub TmA neurons connecting in multiple directions across an LGMD 2 dendrite setting up the anatomical substrate for both short and long range suppression (Cocks PhD thesis, 2020). Acceptance angles of neighbouring columns are 0.7° in the mantis fovea, so 5 pixels subtended 1.2 degrees (Rossel, 1979).

In our experiments, when WFM and target motion are out of phase (100%) or partially out of phase (50% in phase), the WFM generated no overt optomotor behaviour and tracking is supressed in proportion to the spatial distance between target and WFM supporting a role for looming-type motion-detectors in tracking. Prete and Mahaffey, (Prete and Mahaffey, 1993) however, reached a different conclusion: that mantis tracking responses made to movies of walking cricket-like objects showed no evidence of suppression by a nearby WFM and could not be mediated by looming type motion detection, even though the strike did (Prete, 1992; Prete and Mahaffey, 1993; Prete and McLean, 1996). Stimulus presentation in Prete and Mahaffey (Prete and Mahaffey, 1993) was different to the present study, optomotor behaviour was evoked by their background WFM and was in opposition to tracking responses and could have confounded them. Stimuli consisted of a repeated series of five frames that depicted sequential cricket leg positions, as the frames cycled, the crickets appeared to walk across the centre of the screen, in a straight line against stationary or horizontally moving backgrounds (background speed = 2 or 3.5 ° s^-1^). We clearly find that WFM suppresses tracking responses to SF target motion, particularly when the two motions are close, and in phase with one another. Extended motion of the same phase (direction) as the motion of a small central target would occur if the object is too large, and the target is a feature of a much larger object that includes the background and not within the preferred size range for prey (Nityananda et al., 2019; Nityananda et al., 2016).

### Other SFM detectors

In the mantis adding fine detail to a moving target, particularly closely spaced light and dark edges diminishes tracking behaviour. Findings such as these, have been used to indicate spatial antagonism or inhibitory lateral interactions in visual pathways, for example, in small-target motion-detector neurons (STMDs) of male hoverflies where the addition of internal features to a target results in reduced excitation (Nordström et al., 2006), and also in locust looming detectors responses to light and dark edges suppress each other (Pinter, 1979; Rind and Simmons, 1998). In *Drosophila*, inhibitory interactions have been demonstrated directly, in a projection neuron from the lobula, named LC11, which responds to the movement in any direction, of small objects darker than the background, with little or no responses to static flicker, vertically elongated bars, or WFM of gratings, but in which, blocking inhibitory ionic currents eliminates small object sensitivity and instead leads to responses to elongated bars and gratings (Keleş and Frye, 2017; Von Reyn et al., 2017; Wu et al., 2016). In the STMD neurons in dragonflies and male hoverflies, (Bolzon et al., 2009; Nordström, 2012; Wiederman et al., 2008) modelling shows how targets, whose image subtends less than one inter-ommatidial angle, can be visualized against moving clutter, even without relative velocity differences between target and background. The model of the STMD neurons relies on the signature of a tiny dark target moving across a single point in space which is that the tiny target’s leading edge (dark contrast change) is followed a short time later by its trailing edge (bright contrast change). The model incorporates fast adaptation and strong spatial antagonism (lateral inhibition) that leads to the STMD’s sensitivity to the spatiotemporal dynamics associated with tiny target motion. Many STMD receptive fields are much larger than the optimal target sizes, and yet target selectivity is a position-invariant property (Nordström and O’Carroll, 2006). This indicates that in these neurons target size selectivity is unlikely to derive from spatially distinct inhibition within the neuron itself. It is more likely that small-target specificity is generated by “elementary” small-field target-selective subunits located presynaptic to large descending brain output neurons (Wiederman et al., 2008). In mantis tracking the lateral inhibition of SFM by a moving background showed the same spatial tuning as did target tracking itself suggesting that the two manifestations of spatial inhibition could, like STMDs in hoverfies and dragonflies (Bolzon et al., 2009; Nordström, 2012; Nordström et al., 2006; Nordström and O’Carroll, 2006; Wiederman et al., 2008), derive from the same underlying process rather than from any external spatially distinct inhibition. Although the dark target used in our study subtended 14.32 × 14.32° on the eye and was bigger, covering around 580 ommatidia per eye (Rossel, 1979), compared with than that investigated in the hoverfly which covered only 1–3° or 33 ommatidia. The mantises clearly still identified our target as prey rather than predator because they struck at it and never showed defensive behaviour (Yamawaki, 2011).

### Directional responses from looming-type motion-detectors

It is logical that the mechanisms for detecting looming should be similar to that responsible for tracking as there may be a looming component to target motion, particularly as the target comes toward the mantis as tracking brings it into striking range. In response to a looming threat, looming detecting neurons in locusts mediate flight steering manoeuvres and escape jumps that are highly directional (Fotowat et al., 2011; Santer et al., 2006; Santer et al., 2005). In general looming stimuli generate escape and hiding responses directed away from the stimulus (Card and Dickinson, 2008; Hassenstein and Hustert, 1999; Santer et al., 2005). The neuronal substrate that allows insects to steer in a particular direction, based on looming motion, is unknown although simulations show that based on the simultaneously recorded output of left and right LGMD/DCMD neurones in locusts responding to lateral looming objects a robot could be made to steer directionally away from approaching objects (Yue et al., 2010). This bilateral comparison may also be configured to allow steering of a robot towards a target. One candidate brain structure for mediating directional movements in response to looming-type motion would be the central complex (CC)(Heinze and Homberg, 2008; Homberg, 1987). The CC is dominated by visual input and is involved in orientation of an insect in space (Heinze and Homberg, 2008). Neurons of the locust CC are sensitive to looming stimuli, such as expanding dark shapes and to small translating objects (Rosner and Homberg, 2013). Specific response properties of some of the neurons made them candidates for mediating directional behaviours away from or toward approaching objects (Rosner and Homberg, 2013). Several neuron types showed binocular responses to looming objects, and some neurons were excited or inhibited depending on which eye was stimulated. Many of the CC neurons that ramify in the central bridge region for example, determine heading direction and are laid out in order in columns representing different heading directions in space, around 360°. This could allow translation of looming-type responses across the two eyes into a tracking trajectory which brings a target into a range that stereopsis can determine is within striking distance (Nityananda et al., 2018; Rosner et al., 2019).

### Candidate looming-type motion-detecting neurons for mantis target tracking

Investigations exploring the functional organization of the mantis lobula complex in relation to looming-type detection have revealed several large neurones that project from the lobula complex into the protocerebrum. In particular a group of large tangential neurones (T1-3+) that branch in the <u>o</u>uter layers of the lobula complex (O) and then project to the ipsilateral medial or lateral <u>pro</u>tocerebrum (TOpro1-3+ in (Rosner et al., 2020; Rosner et al., 2017); TOproM and TOproL in (Yamawaki, 2019). They are monocular and not involved in stereopsis (Rosner et al., 2020; Rosner et al., 2017) however these neurons not only respond best to SFM of a dark target in any direction over the eyes (Berger, 1985; Rosner et al., 2020; Rosner et al., 2017; Yamawaki, 2019) and the responses of at least one of them (TOproL), are also suppressed by WFM of a striped grating (Yamawaki, 2019). TOproM also gave larger responses to looming verses receding stimuli although its firing rate did not increase as the object approached, suggesting that TOproM was more sensitive to translating objects rather than looming objects but clearly has looming-type sensitivities. Plus both TOpro M and L, like most looming neurons are not in themselves directionally selective for the direction of target translatory motion (Yamawaki, 2019). Potentially their output could be compared on the left and right side to allow directional responses as in the locust (Yue et al., 2010). In summary, our study reveals a possible new role in target tracking for motion pathways featuring long- and short-range suppression, known as lateral inhibition (Barlow, 1961; O’Shea and Rowell, 1975; Pick and Buchner, 1979; Rind, 1987) first characterised in the looming detectors found in insects (Gabbiani et al., 2002; Hatsopoulos et al., 1995; Keleş and Frye, 2017; Rind and Simmons, 1992) then in fish (Temizer et al.), amphibians (Nakagawa and Hongjian, 2010) birds including hummingbirds (Altshuler and Wylie, 2020; Dakin et al., 2016; Goller and Altshuler, 2014; Sun and Frost, 1998), mice (Shang et al., 2015), cats (Liu et al., 2011), and crabs (Oliva and Tomsic, 2014) and potentially represented by the previously described, large TOpro1 or TOproL neurons in the mantis visual system (Yamawaki, 2019).

## Acknowledgements

This research was supported by a BBSRC (Biotechnology and Biological Sciences Research Council) PhD studentship (1219068) to L.J. and it received funding from the European Union’s Horizon 2020 research and innovation programme under the Marie Sklodowska-Curie grant agreement No 691154 STEP2DYNA and agreement No BH295151 LIVCODE to F.C.R. It was also supported by a Leverhulme award Research Leadership Award RL-2012-019 to J.C.A.R. Thanks go to Melissa Bateson for advice and patience on statistical data modelling.

## Contributions

Conceptualization, J.C.A.R., G.T., L.J. and F.C.R.; Methodology, G.T., L.J. and F.C.R.; Software, G.T.; Validation, G.T.; Formal Analysis, G.T.; Investigation, L.J.; Data Curation, L.J.; Resources, F.C.R. and J.C.A.R.; Writing—Original Draft Preparation, L.J.; Writing—Review & Editing, F.C.R.; Visualization, G.T.; Supervision, J.C.A.R. and F.C.R.; Project Administration, J.C.A.R. and F.C.R.; Funding Acquisition, J.C.A.R. and F.C.R.

## Supplementary information

**Movie 1** Demonstration of optomotor responses, filmed from below to wide-field motion of sinusoidally modulated stripes. The responses were scored without the screen being visible to the observer

**Movie 2** Demonstration of tracking responses filmed from below to small-field motion of a dark target moving rapidly across a grey background. A tracking response is generated by the animal with the target tracked each of the 4 times it transits across the screen. The responses were scored without the screen being visible to the observer

